# Efficient gradient-based parameter estimation for dynamic models using qualitative data

**DOI:** 10.1101/2021.02.06.430039

**Authors:** Leonard Schmiester, Daniel Weindl, Jan Hasenauer

## Abstract

**Motivation:** Unknown parameters of dynamical models are commonly estimated from experimental data. However, while various efficient optimization and uncertainty analysis methods have been proposed for quantitative data, methods for qualitative data are rare and suffer from bad scaling and convergence.

**Results:** Here, we propose an efficient and reliable framework for estimating the parameters of ordinary differential equation models from qualitative data. In this framework, we derive a semi-analytical algorithm for gradient calculation of the optimal scaling method developed for qualitative data. This enables the use of efficient gradient-based optimization algorithms. We demonstrate that the use of gradient information improves performance of optimization and uncertainty quantification on several application examples. On average, we achieve a speedup of more than one order of magnitude compared to gradient-free optimization. Additionally, in some examples, the gradient-based approach yields substantially improved objective function values and quality of the fits. Accordingly, the proposed framework substantially improves the parameterization of models from qualitative data.

**Availability:** The proposed approach is implemented in the open-source Python Parameter EStimation TOolbox (pyPESTO). All application examples and code to reproduce this study are available at https://doi.org/10.5281/zenodo.4507613.

## 1 Introduction

Systems biology models based on ordinary differential equations (ODEs) have enabled a profound understanding of many biological processes. Application examples include the study of cellular signal transduction [Bachmann et al., 2011, Adlung et al., 2017] and the prediction of drug treatment outcomes [Hass et al., 2017, Fröhlich et al., 2018]. The ODE models employed in these and other applications commonly comprise parameters, such as reaction rate constants or initial concentrations of biochemical species, which cannot be measured directly and therefore have to be inferred from experimental data [Mitra and Hlavacek, 2019]. This is achieved by optimizing the agreement of the model simulation with experimental data, e.g., by minimizing the sum of squared distances or by maximizing a likelihood function. Various optimization methods have been developed to solve parameter estimation problems. This includes multi-start local optimization methods, global optimization methods and hybrid optimization methods (see Villaverde et al. [2018] for detailed a discussion). Several empirical studies [Schälte et al., 2018, Raue et al., 2013, Villaverde et al., 2018] have shown that optimization methods which use the gradient of the objective function with respect to the parameters tend to be more efficient than gradient-free optimization methods. Yet, while gradient calculation for objective functions using quantitative data is well established [Raue et al., 2013, Fröhlich et al., 2017, Sengupta et al., 2014], respective tools for qualitative data are missing.

A spectrum of experimental setups and techniques provide qualitative observations, meaning that no exact quantitative relation to the concentration e.g. of biochemical species is available [Pargett and Umulis, 2013]. Examples include imaging data for certain stainings [Pargett et al., 2014, Brooks et al., 2012], Förster resonance energy transfer (FRET) data [Birtwistle et al., 2011] or phenotypic observations [Chen et al., 2004]. Although qualitative measurements do not provide numerical values, they contain valuable information to infer parameters [Pargett and Umulis, 2013]. Therefore, several tailored parameter estimation approaches have been developed: (i) Oguz et al. [2013] optimized the number of qualitative observations that were correctly captured by the model. (ii) Mitra et al. used qualitative observations as static penalty functions [Mitra et al., 2018] and proposed a pseudo-likelihood function [Mitra and Hlavacek, 2020]. (iii) Pargett et al. [2014] employed the concept of the optimal scaling approach (introduced by Shepard [1962]), which is based on finding the best possible quantitative representation (so-called surrogate data) of the qualitative observations. This is based on a hierarchical optimization problem, where in an outer optimization loop the model parameters are estimated, and in an inner optimization loop the optimal surrogate data is calculated. The approaches (i)-(iii) facilitate the extraction of information about the model parameters from qualitative data. Yet, the objective functions are either intrinsically discontinuous or an analytical formulation for the objective function gradient was unknown. Accordingly, only gradient-free optimization methods could be employed.

Here, we derive formulas for semi-analytical calculation of the gradients of the objective function arising in the optimal scaling approach. This allows for the use of gradient-based optimization in the outer loop (which optimizes the model parameters), and complements our previous work [Schmiester et al., 2020] on the reformulation of the inner loop (which optimizes the surrogate data). We evaluate our gradient-based framework on several application examples and compare it to gradient-free optimization. We show that the proposed method yields accurate gradients, substantially accelerates parameter estimation and profile calculation and often yields improved final objective function values.

## 2 Method

### 2.1 Mathematical modeling of biological processes

We consider ODE models

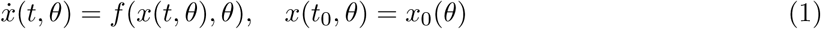

for the dynamics of the concentrations of biochemical species, 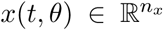. The vector field 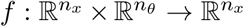 describes the temporal evolution of the modeled species. The unknown model parameters are denoted by 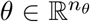 and the initial states at time point *t*_0_ are given by *x*_0_(*θ*). The total numbers of state variables and unknown parameters are denoted by *n*_*x*_ and *n*_*θ*_, respectively.

The state variables *x*(*t, θ*) can be linked to experimental data by introducing an observation function 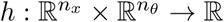, which maps the *n*_*x*_ state variables to an observable, *y*(*t, θ*) ∈ ℝ, via

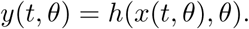

Note that we consider here the case of a single observable to simplify the notation. The general case is captured in the Supplementary Information, Section 2.

**Quantitative data** provide information about the observable *y*(*t, θ*). Yet, the measurements are usually subject to measurement errors, which are often assumed to be i.i.d. additive Gaussian noise.

In this case, the data 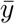 can be linked to the observables by

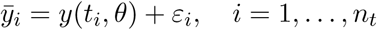

with measurement noise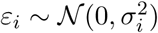, in which *σ*_*i*_ is the standard deviation for the *i*-th observation and *n*_*t*_ denotes the number of time points. Alternative noise models can be used depending on the measurement characteristics (see discussion in Maier et al. [2017]).

**Qualitative data** do not provide information on the values of the observable, but rather on the ordering of different datapoints. Following the notation from Schmiester et al. [2020], we denote a qualitative readout as *z*(*t, θ*) ∈ R. For these readouts no exact quantitative relation to *y*(*t, θ*) is known and only monotonicity in the mapping between *y*(*t, θ*) and *z*(*t, θ*) can be assumed. A qualitative observation provides the ordering of two readouts. It is possible that these observations are indistinguishable (*z*_*i*_ ≈ *z*_*j*_), or that one observable is clearly larger or smaller (*z*_*i*_{*>, <*}*z*_*j*_). We group the different indistinguishable qualitative measurements in *n*_*k*_ different categories which are denoted by 𝒞_*k*_, *k* = 1, …, *n*_*k*_, i.e. *z*_*i*_, *z*_*j*_ ∈ 𝒞_*k*_ ⇒ *z*_*i*_ ≈ *z*_*j*_. We assume in the following that the categories are ordered as 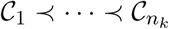.

### 2.2 Parameter estimation using qualitative data

In this study, we build upon the *optimal scaling* approach for parameter estimation using qualitative data [Pargett et al., 2014, Schmiester et al., 2020]. The optimal scaling approach introduces *surrogate data*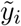, which are the best quantitative representations of the qualitative measurements 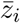.Therefore, for some parameter vector *θ*, the surrogate data 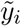 aim to describe the model simulation *y*(*t*_*i*_, *θ*) optimally, while fulfilling the ordering of the qualitative categories (Figure 1A). To this end, we introduce intervals [*l*_*k*_, *u*_*k*_] for each category 𝒞_*k*_, with lower and upper bounds *l*_*k*_ and *u*_*k*_. The intervals are ordered, *u*_*k*_ ≤ *l*_*k*+1_, and the surrogate datapoints can be freely placed within the corresponding interval. As the bounds of the intervals and the surrogate data are a priori unknown, they are subject to optimization. Using a weighted sum of squared distances function, the corresponding optimization problem is

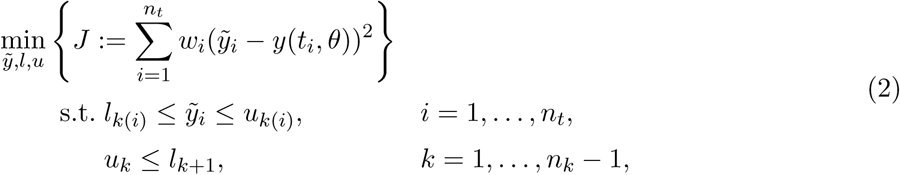

in which *k*(*i*) is the index of the category of the surrogate datapoint 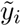 and *w*_*i*_ are datapoint-specific weights. The weights are usually chosen such that the objective function value is independent on the scale of the simulation [Pargett et al., 2014, Schmiester et al., 2020]. The first inequality constraint of (2) guarantees that the surrogate datapoints are placed inside the respective interval, and the second inequality constraint assures that the ordering of the categories is fulfilled.

**Figure 1:**
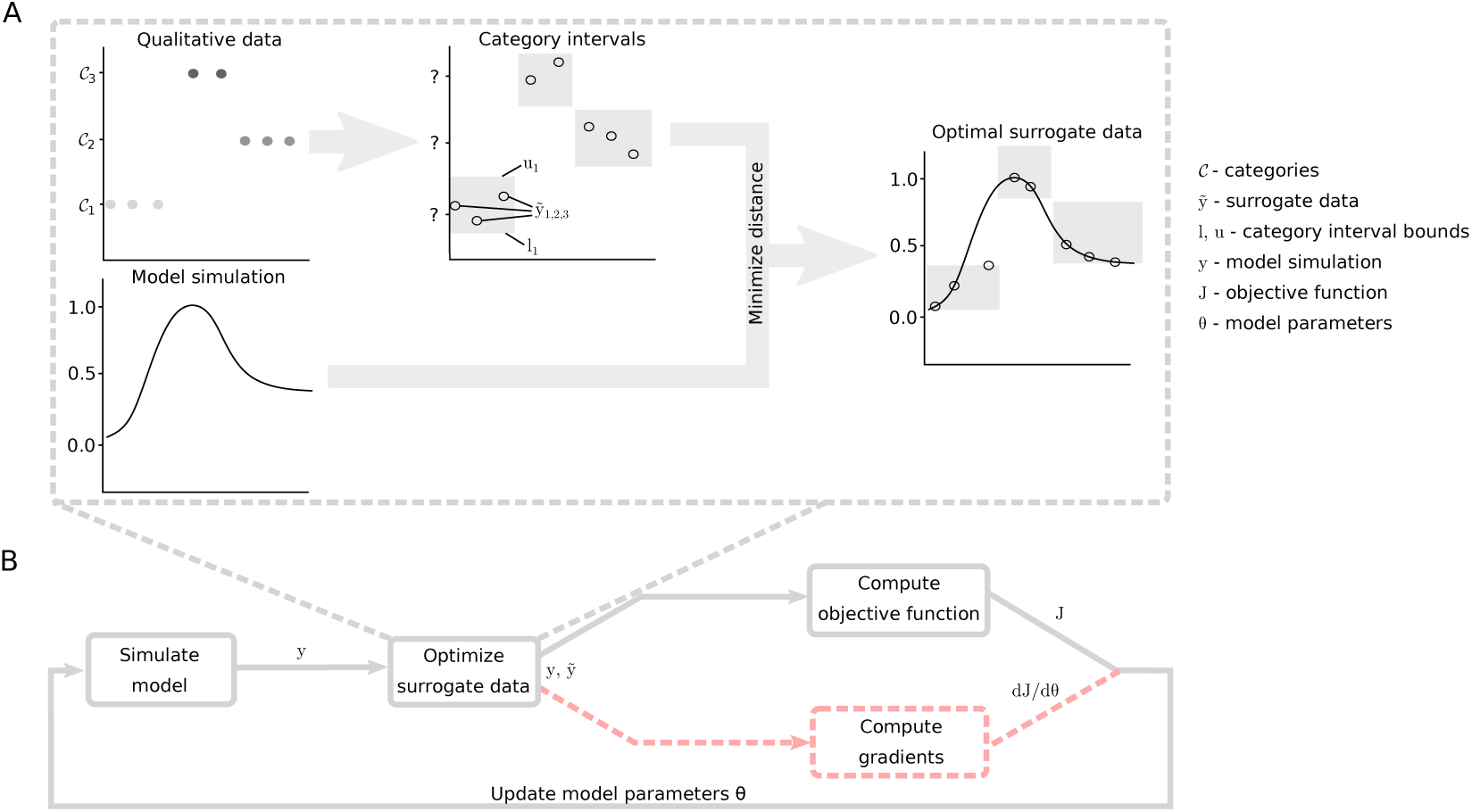
Illustration of the optimal scaling approach. **A**: Qualitative data and model simulation are integrated by introducing intervals for the qualitative categories C, which have to be optimized to minimize the difference to the model simulation. Surrogate data 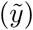 can then be placed optimally inside the intervals [*l, u*]. **B**: Model parameters *θ* are updated iteratively during parameter estimation. For each trial parameter vector, the model output *y* is simulated. Then, the surrogate data is optimized and used to compute the objective function *J* and, if required by the employed optimizer, gradients *dJ/dθ*.

To estimate the unknown model parameters *θ*, the surrogate data optimization (2) can be nested into the model parameter optimization (Figure 1B), yielding the hierarchical optimization problem

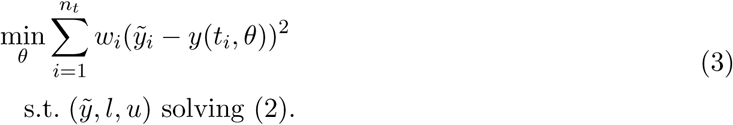

Therefore, in each iteration of the outer optimization (of the model parameters *θ*), the inner constrained optimization problem (2) (for the surrogate data and category bounds) has to be solved. We have previously shown that the inner optimization problem can be simplified to improve efficiency and robustness by only estimating upper bounds *u* and determining *l* and 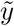 analytically [Schmiester et al., 2020]. However, for the derivation of the gradient computation algorithm introduced in the next section we will consider the full optimization problem (2).

For ease of notation, we rewrite the optimization problem in matrix-vector notation. For this, we denote the collection of all inner optimization variables by 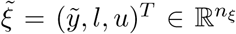 and the vector of simulations by 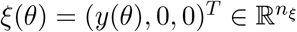which is filled with zeros such that 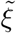 and *ξ* have the same dimension. Here, *n*_*ξ*_ = *n*_*t*_ + 2*n*_*k*_ is the number of inner optimization variables. With this, we can rewrite the optimization problem (3) as a hierarchical problem of the form

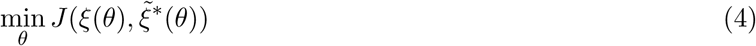

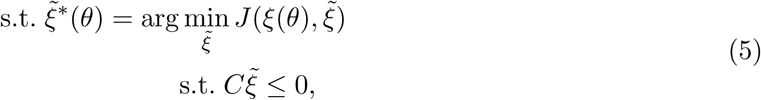

in which 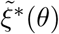 is the optimal value of the inner optimization for a given *θ*. Note that, because the optimal solution to the inner optimization problem (5) depends on the model parameters *θ*, the optimal surrogate data 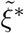 is directly dependent on *θ*. The objective function is given by

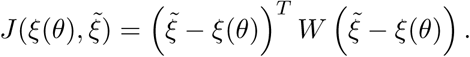

The matrix 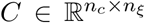 encodes the inequality constraints of the inner problem, with the total number of constraints *n*_*c*_, and the weight matrix *W* is given by

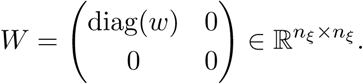

*W* is augmented with zeros such that the bounds *l* and *u* do not contribute to the objective function *J* and the dimensions of *W* and *C* are consistent, which is necessary for the following calculations. An illustrating example of the reformulation is given in the Supplementary Information, Section 3.

### 2.3 Gradient computation for the optimal scaling objective function

In this section, we derive an algorithm to calculate the gradients of the optimal scaling objective function, based on ideas from the field of bi-level optimization [Kolstad and Lasdon, 1990, Fiacco, 1976]. We are interested in calculating the derivative of the objective function *J* with respect to the model parameters *θ*, evaluated at the optimal surrogate data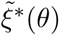, which is given by

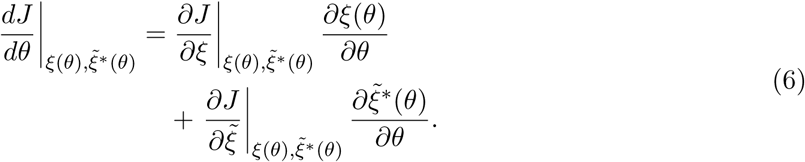

All parts of (6) except for 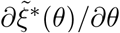 can be easily calculated (see Supplementary Information, Section 1 for details). To obtain 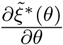, we introduce the Lagrangian function 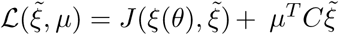, with Lagrange multipliers 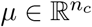. The necessary first order optimality conditions of problem for a given *θ* are then given by the Karush-Kuhn-Tucker conditions [Boyd and Vandenberghe, 2004]

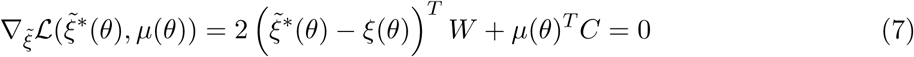

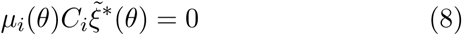

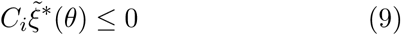

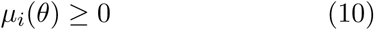

for *i* = 1, …, *n*_*c*_, where *C*_*i*_ is the i-th row of *C*. Given the optimal values of the inner optimization variables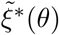, the Lagrange multiplier *µ* can be calculated by solving (7)–(10). To obtain 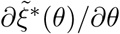, we calculate the derivative of the optimality conditions (7) and (8) w.r.t. *θ*_*j*_, which results in

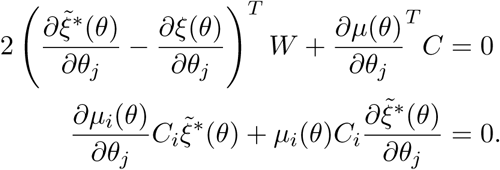

This yields a linear system of equations that needs to be solved for every parameter *θ*_*j*_:

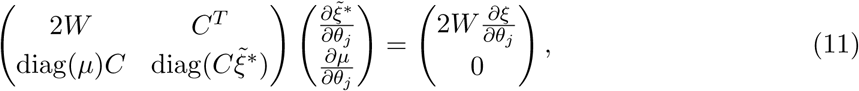

where we omitted the dependency of *µ, ξ* and 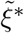 on *θ* for simplicity of presentation. After solving (11), we can calculate the gradients of the objective function w.r.t. the model parameters *θ* using equation (6). To summarize, the gradient computation scheme consists of the following steps:

1. Simulate the ODE (1) to obtain *ξ*(*θ*) an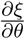.
2. Calculate optimal surrogate data 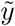 and category bounds *l, u* by solving (2) or a reformulation of this problem [Schmiester et al., 2020].
3. Solve the optimality conditions (7) - (10) for the Lagrange multiplier *µ*.
4. Solve the linear system of equations (11) to obtain 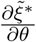.
5. Evaluate the gradient 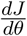 of the objective function using equation (6).

In practice, it is sometimes preferable to choose weights, that are dependent on the parameters *θ* or the simulation *ξ*. Additionally, minimal sizes on the intervals *s*(*θ, ξ*(*θ*)) ≤ *u*_*k*_ − *l*_*k*_ and on the gaps between the intervals *g*(*θ, ξ*(*θ*)) ≤ *l*_*k*+1_ − *u*_*k*_, which can also depend on *θ* and *ξ*, can be imposed. Assuming that *g*(*θ, ξ*(*θ*)), *s*(*θ, ξ*(*θ*)) and *W* (*θ, ξ*(*θ*)) are differentiable functions, similar formulas for gradient computation can be derived (see Supplementary Information, Section 1). Collecting minimal interval gaps *g* and interval sizes *s* in the vector *d*, yielding the inequality constraints 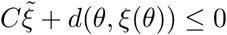, we obtain the linear system of equations

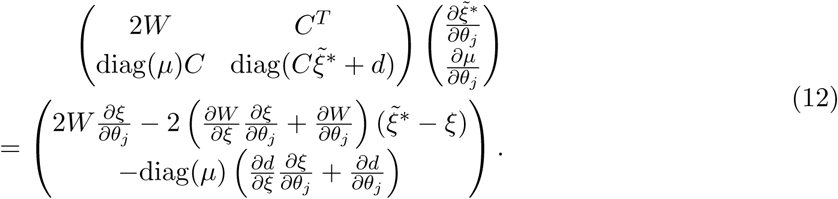

The linear systems (11) and (12) are sparse and can be solved efficiently.

### 2.4 Implementation

We implemented the gradient calculation method in the Python Parameter EStimation TOolbox (pyPESTO) [Schälte et al., 2020]. The qualitative data can be defined using an extension to the PEtab format, which is a standardized format for the definition of parameter estimation problems [Schmiester et al., 2021]. Model simulation was carried out using the AMICI toolbox [Fröhlich et al., 2020]. Parameter estimation was performed using multi-start local optimization with 500 starts per model and method. To guarantee comparability, optimizations were started from the same initial parameters for each method. For gradient-free optimization we used the Powell algorithm implemented in the SciPy package [Jones et al., 2001], which performed well among SciPy’s gradient-free optimizers in a previous study using the optimal scaling approach [Schmiester et al., 2020]. For gradient-based optimization we used SciPy’s L-BFGS-B algorithm, which is the default optimizer in pyPESTO. For more details on the implementation, we refer to the Supplementary Information, Section 4.

## 3 Results

### 3.1 Model overview

To analyze the gradient algorithm and compare it to gradient-free optimization, we considered six models. We included one toy model and five larger application examples. An overview of the models and datasets used for parameter estimation is given in Table 1.

**Table 1:**
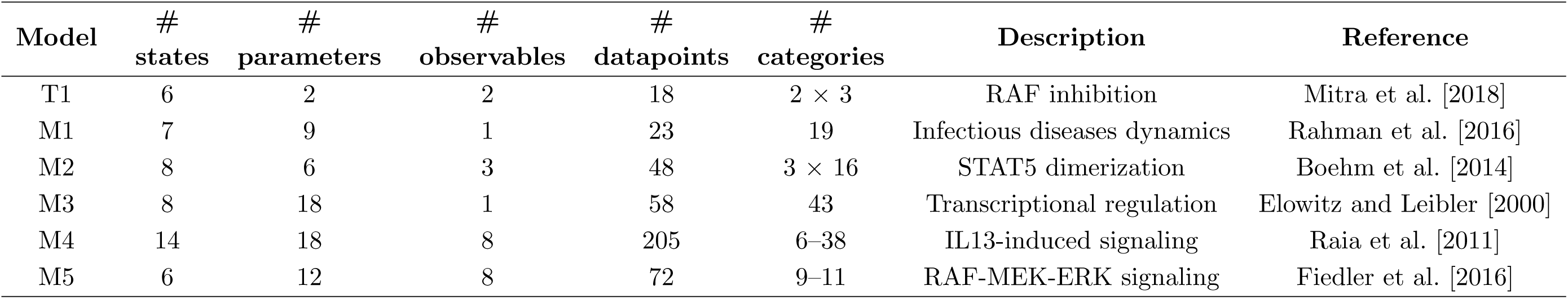
Key numbers of the different considered models and datasets used for parameter estimation.

T1 is a small model used for illustration. Models M1-M5 are published models with experimental data and describe different biological processes. They are taken from a collection of parameter estimation problems in the PEtab format, which is based on the benchmark collection by Hass et al. [2019]. The models comprise different numbers of datapoints and unknown parameters and were originally calibrated on quantitative measurements. These quantitative measurements were converted to qualitative observations based on their ordering, where we assumed that measurements are indistinguishable, if their numeric values are equal. Additionally, we assumed that data was comparable within an observable but not across observables, leading to one optimal scaling problem per observable that needed to be solved.

### 3.2 Semi-analytical approach yields accurate gradients

As the gradient computation algorithm involves numerically solving a linear system of equations, we first evaluated the accuracy of the obtained solution. We considered the model T1 and compared the semi-analytical gradients with gradients obtained using a finite difference approach at different parameter vectors (Figure 2A). The analysis revealed that for all tested parameters the approaches yielded almost identical gradients (with absolute differences *<* 10^*−*9^). We additionally compared computation times of finite differences and our semi-analytical gradient algorithm for models M1–M5, which showed that finite differences require substantially more computation time (Supplementary Information, Figure S1). Therefore, we restrict ourselves to gradient computations using the semi-analytical approach in the following analysis.

**Figure 2:**
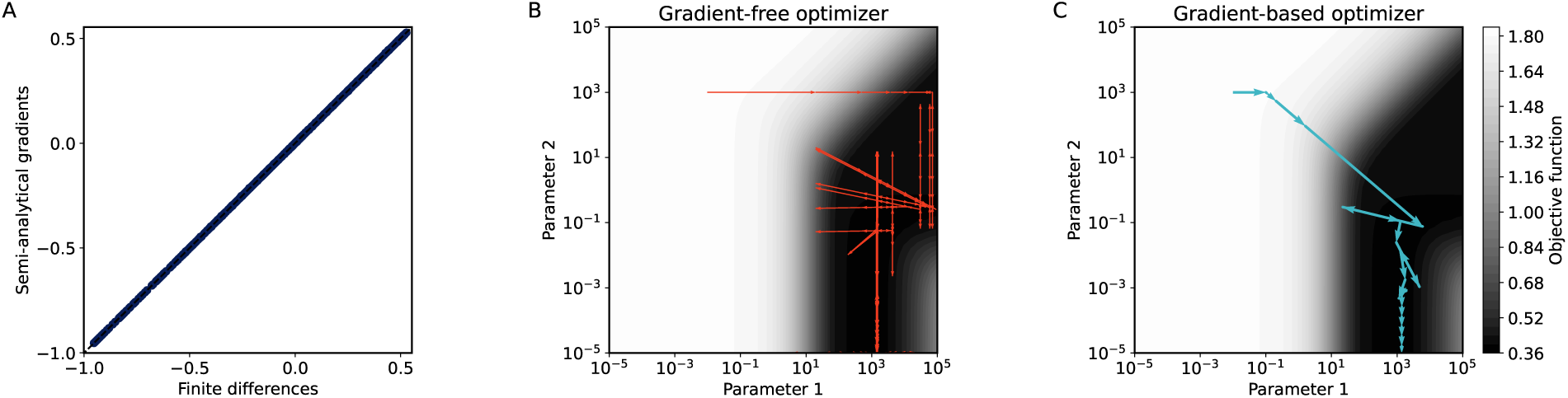
**A:** Absolute gradients of the two parameters of model T1 evaluated at 2500 uniformly sampled parameter vectors using the semi-analytical approach and central finite differences. **B & C:** Objective function landscape and optimizer trajectories of a gradient-free (B) and a gradient-based (C) optimizer.

### 3.3 Gradient information increases optimizer efficiency

To illustrate the differences of gradient-free and gradient-based optimization, we estimated parameters for both optimizers starting from the same initial parameters. As the model T1 only contains two parameters, we inspected the whole objective function landscape and the respective optimizer trajectories (Figure 2B & C). While the gradient-free optimizer used a rather naive updating scheme (Figure 2B) that often moved along sub-optimal directions, the gradient-based optimizer moved towards the optimal point within a few iterations (Figure 2C), indicating a potential advantage of using gradient-based optimizers.

To assess the performance of gradient-based and gradient-free optimization in a realistic setting, we performed multi-start local optimization for the application examples M1–M5. The results show considerably reduced computation times of the gradient-based optimization for all models (Figure 3A). Depending on the considered model, the median CPU times are reduced by 1–2 orders of magnitude.

**Figure 3:**
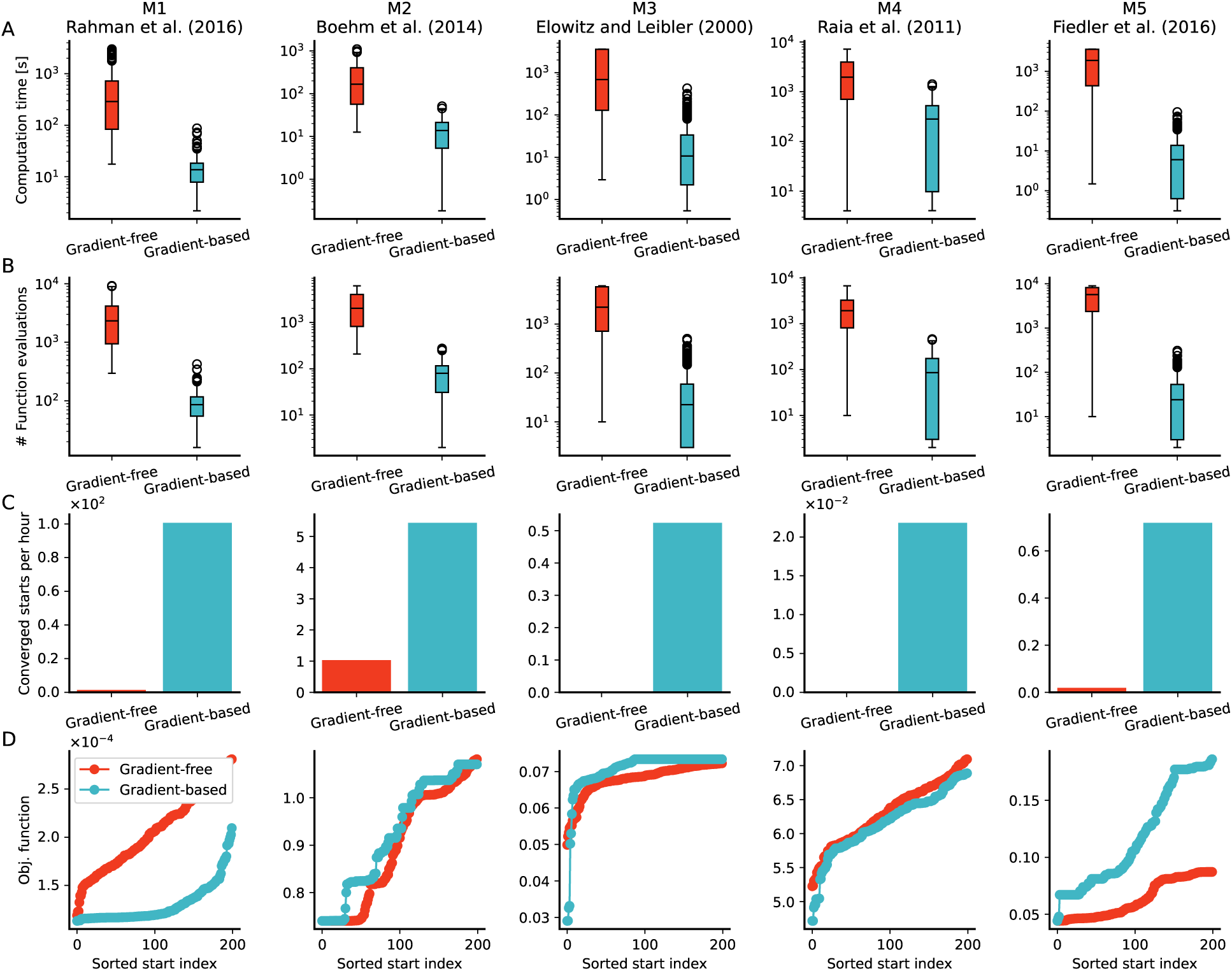
Optimization results for all models using gradient-free and gradient-based optimization for 500 local optimizations. **A:** Computation times until the optimizer terminates per local optimization. **B:** Number of function evaluations per local optimization. **C:** Converged starts per hour. A start is considered converged, if the absolute difference to the overall best value is less than 10^*−*4^ (see Supplementary Information, Figure S2 for results using different thresholds). **D:** Waterfall plots for all 5 considered models using gradient-free and gradient-based optimization. Best 200 starts out of a total of 500 are shown. See Supplementary Information, Figure S3 for results of all starts.

As illustrated in Figure 2, the gradient-based optimizer used a more intelligent updating scheme during optimization, resulting in reduced numbers of objective function evaluations. The reduction in function evaluations could also be observed for all employed models (Figure 3B). Even though a single evaluation is more costly when gradients need to be calculated (Supplementary Information, Figure S1), the reduced number of necessary evaluations outweighs this, explaining the improved computation times.

In addition to the computation times, we also incorporated the final objective function values into our analysis by considering the number of local optimization runs which achieved values close to the overall best value (Figure 3C). This revealed substantially improved efficiency of the gradient-based optimizer for all models. This result could be observed independent of the threshold for convergence to the optimal point (Supplementary Information, Figure S2). The improved efficiency of the gradient-based optimizer could be observed even for the models M2 and M5, for which the gradient-free optimizer found the optimal value more often (Figure 3D). A possible explanation for the sometimes larger number of converged starts using gradient-free optimization could be flat regions in the objective function which lead to a termination of the gradient-based optimization.

### 3.4 Gradient-based optimization yields improved model fits

The waterfall plots show that the gradient-based optimization yielded equal (M1, M2 and M5) or even better (M3 and M4) final objective function values compared to gradient-free optimization (Figure 3D and Supplementary Information, Figure S3). We simulated the models for which larger differences in the best obtained objective function values were observed to analyze if the different parameters resulted in substantial differences in the model fits. Especially for model M3 we observed a better representation of the data with the parameters from gradient-based optimization (Figure 4). Indeed, only the parameters obtained using the gradient-based optimization correctly captured the oscillations of the measurements.

**Figure 4:**
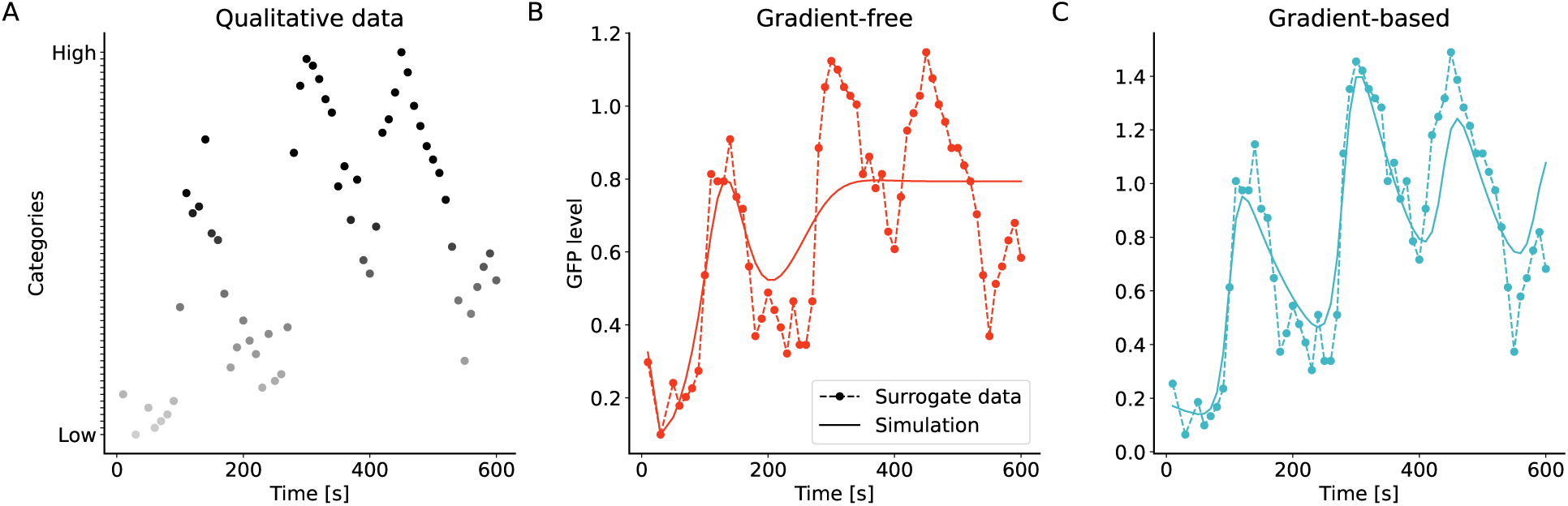
Fits of model and qualitative data for model M3. **A:** Qualitative observations. The gray-scale indicates the ordering of the qualitative datapoints. **B & C:** Simulated data and optimal surrogate data for the best parameters from gradient-free (B) and gradient-based (C) optimization. The surrogate data are ordered according to the qualitative data (A), but can have different numeric values in B and C as they are optimized to the respective model simulation.

For the model M4, we also observed smaller improvements in the model fits for some observables, when using the parameters obtained from gradient-based optimization (Supplementary Information, Figure S4).

### 3.5 Gradient-based approach facilitates uncertainty quantification

Qualitative data is often considered to be less informative than quantitative measurements. As this can result in reduced parameter identifiability, it is even more important to assess the uncertainties associated with the estimated parameters when using qualitative measurements. For uncertainty analysis, we used objective function ratio profiles analogously to profile likelihoods in the case of a likelihood function [Raue et al., 2009]. While the objective function differences in the profiles cannot easily be interpreted statistically, they can still be valuable to indicate uncertainties of the estimated parameters. As a proof of concept, we calculated objective function profiles for the model M2 (Figure 5A). The gradient-based approach yielded mostly smooth profiles indicating that several parameters could be well identified using the qualitative dataset. In contrast, the gradient-free approach resulted in several discontinuities in the profiles probably caused by impaired optimization. This shows that only the gradient-based approach was able to yield meaningful profiles for this model. In addition to the improved profiles, the gradient-based approach required on average an order of magnitude less computation time than the gradient-free optimizer (Figure 5B).

**Figure 5:**
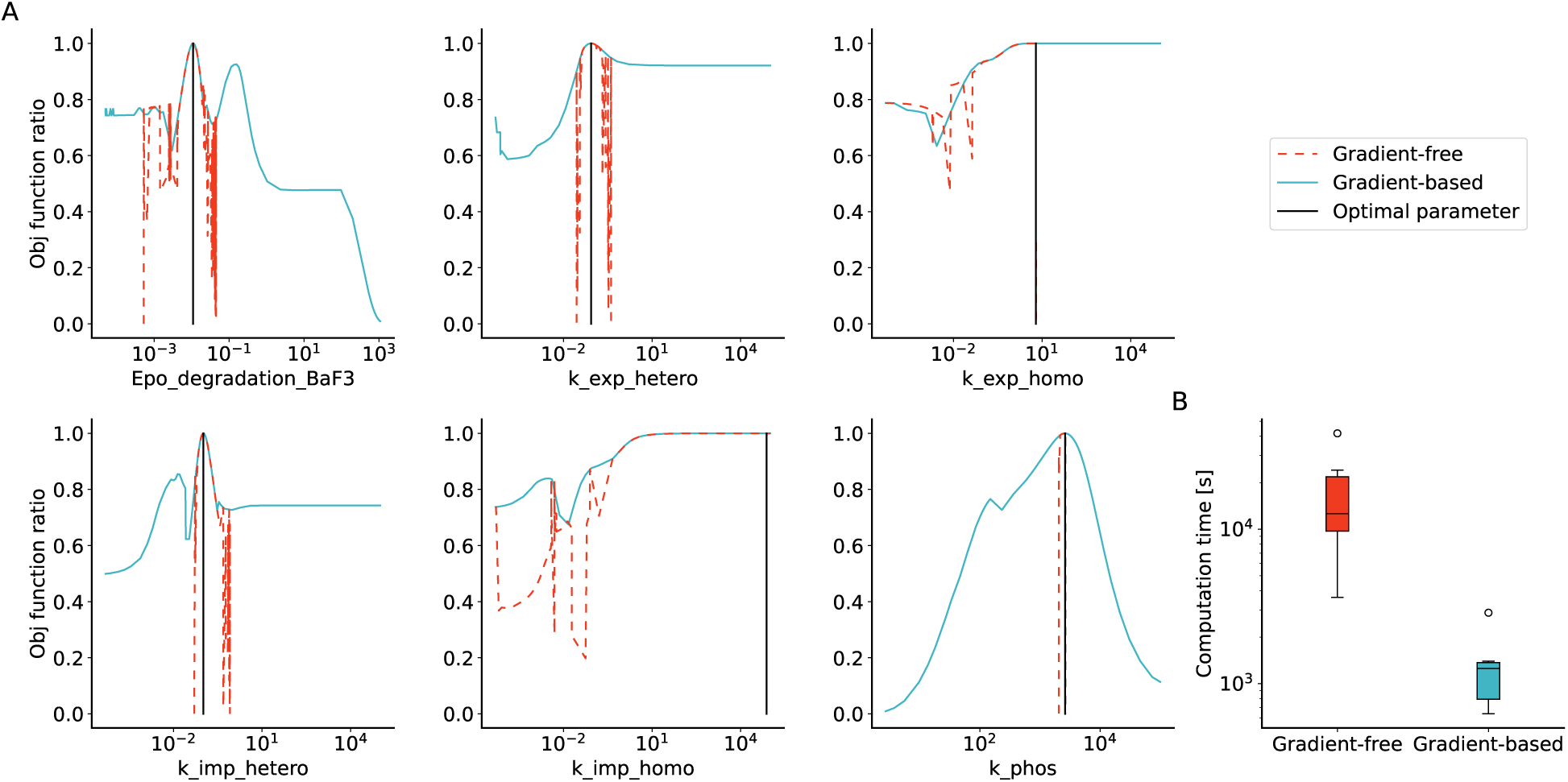
**A:** Objective function ratio profiles for the six parameters of model M2 using gradient-free and gradient-based optimization. Shown is the ratio of the overall best objective function value and the value along the profile. Profiles were initiated on the overall best found parameter. **B:** Boxplots for computation times for the profiles per parameter.

## 4 Discussion

Qualitative data can contain valuable information for parameter estimation but current methods to integrate such data are computationally demanding and more efficient algorithms are required. Here, we developed a framework for semi-analytical computation of gradients for the optimal scaling approach. We validated the accuracy of the obtained gradients by comparing them to finite differences and assessed the advantage of using gradient information on five application examples by performing optimization with a gradient-free and a gradient-based algorithm. This revealed speedups of more than one order of magnitude using the gradient-based approach. Additionally, the gradient-based algorithm resulted in equal or even improved final objective function values and model fits. The gradient-based approach was further used to reliably calculate objective function profiles to assess the uncertainty of parameter estimates when using qualitative data.

A linear system needs to be solved for every parameter to obtain the gradients of the optimization problem. As this is the most time consuming part during gradient computation, more efficient approaches could further decrease computation times. The computation time could for instance be reduced by splitting it into active and inactive constraints [Kolstad and Lasdon, 1990] or by parallelizing gradient computation over the parameters. Complementary to this, it remains open whether – similar to other hierarchical optimization approaches [Schmiester et al., 2019] – adjoint sensitivity analysis can be used to further accelerate optimization [Fröhlich et al., 2017]. Another possible extension would be the derivation of second-order derivatives that could be used for parameter estimation and profile calculation [Stapor et al., 2018].

It will often be advantageous to combine qualitative and quantitative measurements. This can in principle be done by formulating a similar objective function for quantitative data by replacing the surrogate data with the measured quantitative values. As the gradient for quantitative data can easily be calculated, an overall objective function value and gradient can be obtained by summing up the respective values for qualitative and quantitative data.

In conclusion, we developed a framework to compute gradient information for parameter estimation problems that include qualitative data and showed that it substantially improves computational efficiency. The open-source implementation of the approach we provide will facilitate reusability and might improve the usage of qualitative data for the parameterization of quantitative models.

## 6 Author Contributions

L.S. and J.H. derived the theoretical foundation. L.S. wrote the implementations and performed the case study. L.S., J.H. and D.W. analyzed the results. All authors wrote and approved the final manuscript.

