## Supplementary Information for "Efficient gradient-based parameter estimation for dynamic models using qualitative data"

#### List of Figures

S1 Computation times for an objective function evaluation for 50 randomly sampled parameter vectors for all five application examples . . . . . . . . . . . . 7

S2 Number of converged starts per hour for different convergence thresholds . . 8

S3 Waterfall plots for all models using gradient-free and gradient-based optimization 9

S4 Model simulation and optimal surrogate data for the best found parameters for model M4 . . . . . . . . . . . . . . . . . . . . . . . . . . . . . . . . 10

#### 1 Derivation of gradient formulas

Here, we derive the algorithm for calculating the gradient of the optimal scaling objective function in detail for the case of one observable. First, we briefly recapitulate the necessary notation and prerequisites. We consider the general case of parameter and simulation dependent weights $W(\theta, \xi(\theta))$ and minimal interval and gap sizes which are collected in the vector $d(\theta, \xi(\theta))$. The parameters of the inner optimization problem, $\tilde{y}$, $l$ and $u$, are collected in the parameter vector $\tilde{\xi} = (\tilde{y}, l, u)^T$. The simulations are collected in $\xi(\theta) = (y(\theta), 0, 0)^T$. The optimization problem is given by

$$\min_{\theta} J(\theta, \xi(\theta), \tilde{\xi}^*(\theta)) \tag{1}$$

$$\begin{aligned} \text{s.t. } \tilde{\xi}^*(\theta) = \arg\min_{\tilde{\xi}} & \, J(\theta, \xi(\theta), \tilde{\xi}) \\ \text{s.t. } & \, C\tilde{\xi} + d(\theta, \xi(\theta)) \leq 0, \end{aligned} \tag{2}$$

with the objective function

$$J(\theta, \xi(\theta), \tilde{\xi}) = \left(\tilde{\xi} - \xi(\theta)\right)^T W(\theta, \xi(\theta)) \left(\tilde{\xi} - \xi(\theta)\right). \tag{3}$$

$\tilde{\xi}^*(\theta)$ are the optimal surrogate data and interval bounds, which solve the problem (2). Note that while $\tilde{\xi}^*(\theta)$ depends directly on the parameters $\theta$, for $\tilde{\xi}$ this depends on whether the inner or outer problem is considered. In the inner problem $\tilde{\xi}$ does not depend on $\theta$, yet, the optimal solution $\tilde{\xi}^*(\theta)$ of the inner problem (which is used in the outer problem) depends on $\theta$. We are now interested in calculating the derivatives of $J$ w.r.t. the outer parameters $\theta$ evaluated at $\tilde{\xi}^*(\theta)$. This can be calculated by

$$\left.\frac{dJ}{d\theta}\right|_{\theta,\xi(\theta),\tilde{\xi}^*(\theta)} = \left.\frac{\partial J}{\partial \theta}\right|_{\theta,\xi(\theta),\tilde{\xi}^*(\theta)} + \left.\frac{\partial J}{\partial \xi}\right|_{\theta,\xi(\theta),\tilde{\xi}^*(\theta)} \frac{\partial \xi(\theta)}{\partial \theta} + \left.\frac{\partial J}{\partial \tilde{\xi}}\right|_{\theta,\xi(\theta),\tilde{\xi}^*(\theta)} \frac{\partial \tilde{\xi}^*(\theta)}{\partial \theta}. \tag{4}$$

The partial derivatives of $J$ are

$$\left.\frac{\partial J}{\partial \theta}\right|_{\theta,\xi(\theta),\tilde{\xi}^*(\theta)} = (\tilde{\xi}^*(\theta) - \xi(\theta))^T \left.\frac{\partial W}{\partial \theta_j}\right|_{\theta,\xi(\theta)} (\tilde{\xi}^*(\theta) - \xi(\theta)), \tag{5}$$

$$\left.\frac{\partial J}{\partial \xi}\right|_{\theta,\xi(\theta),\tilde{\xi}^*(\theta)} = -2(\tilde{\xi}^*(\theta) - \xi(\theta))^T W(\theta, \xi(\theta)) + (\tilde{\xi}^*(\theta) - \xi(\theta))^T \left.\frac{\partial W}{\partial \xi}\right|_{\theta,\xi(\theta)} (\tilde{\xi}^*(\theta) - \xi(\theta)), \tag{6}$$

$$\left.\frac{\partial J}{\partial \tilde{\xi}}\right|_{\theta,\xi(\theta),\tilde{\xi}^*(\theta)} = 2(\tilde{\xi}^*(\theta) - \xi(\theta))^T W(\theta, \xi(\theta)). \tag{7}$$

The evaluation of these partial derivatives requires the optimal surrogate data $\tilde{\xi}^*(\theta)$ and simulated observable $\xi(\theta)$. Depending on the structure of $W$, also the sensitivities of the simulated observable $\frac{\partial \xi(\theta)}{\partial \theta}$ is required. These components are accessible as solutions of an optimization problem, an ODE, or a forward sensitivity equation.

For the evaluation of the objective function gradient, we additionally need to calculate the derivative of the optimal surrogate data $\tilde{\xi}^*$ w.r.t. the parameters $\theta$. This is the derivative of an optimal solution. To assess it, we follow the ideas from Fiacco (1976) and calculate the derivatives of the first order optimality conditions of the inner optimization problem (2) w.r.t. $\theta$. With the Lagrangian function

$$\mathcal{L}(\tilde{\xi}, \mu) = J(\theta, \xi(\theta), \tilde{\xi}) + \mu^T (C\tilde{\xi} + d(\theta, \xi(\theta))), \tag{8}$$

and Lagrange multipliers $\mu \in \mathbb{R}^{n_c}$, the necessary first order optimality conditions of problem (2) are

$$\nabla_{\tilde{\xi}} \mathcal{L}(\tilde{\xi}^*(\theta), \mu) = 2(\tilde{\xi}^*(\theta) - \xi(\theta))^T W(\theta, \xi(\theta)) + \mu(\theta)^T C = 0 \tag{9}$$

$$\mu_i(\theta)(C_i \tilde{\xi}^*(\theta) + d_i(\theta, \xi(\theta))) = 0, \quad \text{for } i = 1, \ldots, n_c \tag{10}$$

$$C_i \tilde{\xi}^*(\theta) + d_i(\theta, \xi(\theta)) \leq 0, \quad \text{for } i = 1, \ldots, n_c \tag{11}$$

$$\mu_i(\theta) \geq 0, \quad \text{for } i = 1, \ldots, n_c. \tag{12}$$

For a convex objective function, which we have in this case (Schmiester *et al.*, 2020), these conditions are necessary and sufficient for an optimum. To obtain the desired derivatives $\frac{\partial \tilde{\xi}^*}{\partial \theta}$, we calculate the derivatives of equations (9) and (10) w.r.t. $\theta_j$:

$$2\left(\frac{\partial \tilde{\xi}^*(\theta)}{\partial \theta_j} - \frac{\partial \xi(\theta)}{\partial \theta_j}\right)^T W(\theta, \xi(\theta)) + 2\left(\tilde{\xi}^*(\theta) - \xi(\theta)\right)^T \left(\frac{\partial W(\theta, \xi(\theta))}{\partial \theta_j} + \frac{\partial W(\theta, \xi(\theta))}{\partial \xi}\frac{\partial \xi}{\partial \theta_j}\right) + \frac{\partial \mu(\theta)^T}{\partial \theta_j} C = 0 \tag{13}$$

$$\frac{\partial \mu_i(\theta)}{\partial \theta_j}\left(C_i \tilde{\xi}^*(\theta) + d_i(\theta, \xi(\theta))\right) + \mu_i(\theta)\left(C_i \frac{\partial \tilde{\xi}^*(\theta)}{\partial \theta_j} - \left(\frac{\partial d(\theta, \xi(\theta))}{\partial \theta_j} + \frac{\partial d(\theta, \xi(\theta))}{\partial \xi}\frac{\partial \xi(\theta)}{\partial \theta_j}\right)\right) = 0. \tag{14}$$

This yields the following linear system of equations that can be solved for $\frac{\partial \tilde{\xi}^*(\theta)}{\partial \theta}$ and $\frac{\partial \mu(\theta)}{\partial \theta}$ for each parameter $\theta_j$:

$$\begin{pmatrix} 2W(\theta, \xi(\theta)) & C^T \\ \operatorname{diag}(\mu(\theta))C & \operatorname{diag}(C\tilde{\xi}^*(\theta) + d(\theta, \xi(\theta))) \end{pmatrix} \begin{pmatrix} \frac{\partial \tilde{\xi}^*(\theta)}{\partial \theta_j} \\ \frac{\partial \mu(\theta)}{\partial \theta_j} \end{pmatrix} = \begin{pmatrix} 2W(\theta, \xi(\theta))\frac{\partial \xi(\theta)}{\partial \theta_j} - 2\left(\frac{\partial W(\theta, \xi(\theta))}{\partial \theta_j} + \frac{\partial W(\theta, \xi(\theta))}{\partial \xi}\frac{\partial \xi}{\partial \theta_j}\right)(\tilde{\xi}^*(\theta) - \xi(\theta)) \\ -\operatorname{diag}(\mu(\theta))\left(\frac{\partial d(\theta, \xi(\theta))}{\partial \theta_j} + \frac{\partial d(\theta, \xi(\theta))}{\partial \xi}\frac{\partial \xi(\theta)}{\partial \theta_j}\right) \end{pmatrix}. \tag{15}$$

After solving this linear system, we can calculate the gradients of the objective function $J$ w.r.t. the model parameters $\theta$.

Special case of constant $W$ and $d = 0$: The gradients for the special case of a weight matrix which is independent of the parameters and the model observables, and zero minimal interval and gap sizes, i.e. $d = 0$ and $W(\theta, \xi) = W$, can be easily derived from the general result stated above. The linear system determining the derivative of the optimal solution is given by

$$\begin{pmatrix} 2W & C^T \\ \operatorname{diag}(\mu(\theta))C & \operatorname{diag}(C\tilde{\xi}^*(\theta)) \end{pmatrix} \begin{pmatrix} \frac{\partial \tilde{\xi}^*(\theta)}{\partial \theta_j} \\ \frac{\partial \mu}{\partial \theta_j} \end{pmatrix} = \begin{pmatrix} 2W\frac{\partial \xi(\theta)}{\partial \theta_j} \\ 0 \end{pmatrix}. \tag{16}$$

Additionally, $\frac{\partial J}{\partial \theta}\big|_{\theta,\xi(\theta),\tilde{\xi}^*(\theta)}$ vanishes and $\frac{\partial J}{\partial \xi}\Big|_{\theta,\xi(\theta),\tilde{\xi}^*(\theta)}$ reduces to

$$\left.\frac{\partial J}{\partial \xi}\right|_{\theta,\xi(\theta),\tilde{\xi}^*(\theta)} = -2(\tilde{\xi}^*(\theta) - \xi(\theta))^T W. \tag{17}$$

### 2 Optimal scaling approach for multiple observables

In this section, we extend the case of a single observable presented in the main manuscript to the more general case of $n_y$ observables, i.e. $y(t,\theta) \in \mathbb{R}^{n_y}$. Quantitative data is then linked to observables via

$$\bar{y}_{m,i} = y_m(t_i,\theta) + \varepsilon_{m,i}, \quad m = 1,\ldots,n_y, \quad i = 1,\ldots,n_t. \tag{18}$$

For qualitative data, we assume that the ordering of datapoints within one observable is known, but no information on the relation of datapoints across different observables is available. We denote this by introducing a group of $n_{k^m}$ categories for each observable $y_m$, i.e. $\mathcal{C}_{k^m}$, $k^m = 1,\ldots,n_{k^m}$. The respective intervals for the categories are denoted by $[l_{k^m}, u_{k^m}]$. We assume that the categories are ordered as $\mathcal{C}_1 \prec \ldots, \mathcal{C}_{n_{k^m}}$ and that no relation between categories $\mathcal{C}_{k^m}, \mathcal{C}_{k^{m'}}$ are known for $m' \neq m$. The optimal scaling approach then consists of $n_y$ inner subproblems, which need to be solved:

$$\begin{aligned} &\min_{\tilde{\xi}_m} \left\{ J_m := \left(\tilde{\xi}_m - \xi_m(\theta)\right) W_m \left(\tilde{\xi}_m - \xi_m(\theta)\right) \right\} \\ &\quad \text{s.t. } C_m \tilde{\xi}_m \leq 0 \end{aligned} \tag{19}$$

for $m = 1,\ldots,n_y$. Here, $\tilde{\xi}_m$ is the vector of the surrogate data $\tilde{y}_m$ and category bounds $l_m, u_m$, and $\xi_m(\theta)$ the vector of simulations $y_m(t,\theta)$, belonging to the observable with index $m$. $C_m$ contains the constraints of the categories of this observable. The overall objective function value can be calculated by summing over the $n_y$ values obtained from solving (19). Therefore, the optimization problem is given by

$$\min_{\theta} \sum_{m=1}^{n_y} J_m(\theta, \tilde{\xi}_m^*(\theta)) \tag{20}$$

$$\text{s.t. } \left\{ \begin{aligned} \tilde{\xi}_m^*(\theta) = \arg\min_{\tilde{\xi}_m} J_m(\theta, \tilde{\xi}_m) \\ \text{s.t. } \; C_m \tilde{\xi}_m \leq 0 \end{aligned} \right\} \quad \forall m = 1,\ldots,n_y \tag{21}$$

Similarly, the gradient can be calculated via

$$\sum_{m=1}^{n_y} \left.\frac{dJ_m}{d\theta}\right|_{\theta,\xi_m(\theta),\tilde{\xi}_m(\theta)} \tag{22}$$

with

$$\left.\frac{dJ_m}{d\theta}\right|_{\theta,\xi_m(\theta),\tilde{\xi}_m(\theta)} = \left.\frac{\partial J_m}{\partial \theta}\right|_{\theta,\xi_m(\theta),\tilde{\xi}^*_m(\theta)} + \left.\frac{\partial J_m}{\partial \xi_m}\right|_{\theta,\xi_m(\theta),\tilde{\xi}^*_m(\theta)} \frac{\partial \xi_m(\theta)}{\partial \theta} + \left.\frac{\partial J_m}{\partial \tilde{\xi}_m}\right|_{\theta,\xi_m(\theta),\tilde{\xi}^*_m(\theta)} \frac{\partial \tilde{\xi}^*_m(\theta)}{\partial \theta} \tag{23}$$

### 3 Reformulation example

To illustrate the reformulation of the optimal scaling problem in matrix-vector notation, we consider a minimal example of two readouts from two categories $z_1 \in \mathcal{C}_1$, $z_2 \in \mathcal{C}_2$ at timepoints $t_1, t_2$, with $\mathcal{C}_1 \prec \mathcal{C}_2$. The inner optimization problem of the optimal scaling method for this example is

$$\min_{\tilde{y}_1,\tilde{y}_2,l_1,l_2,u_1,u_2} w_1(\tilde{y}_1 - y(t_1,\theta))^2 + w_2(\tilde{y}_2 - y(t_2,\theta))^2 \tag{24}$$

$$\text{s.t. } l_1 \leq \tilde{y}_1 \leq u_1 \tag{25}$$

$$l_2 \leq \tilde{y}_2 \leq u_2 \tag{26}$$

$$u_1 \leq l_2. \tag{27}$$

We can reformulate the objective function in the form (3) using

$$\tilde{\xi} = \begin{pmatrix} \tilde{y}_1 \\ \tilde{y}_2 \\ l_1 \\ l_2 \\ u_1 \\ u_2 \end{pmatrix}, \xi(\theta) = \begin{pmatrix} y(t_1,\theta) \\ y(t_2,\theta) \\ 0 \\ 0 \\ 0 \\ 0 \end{pmatrix}, W = \begin{pmatrix} w_1 & 0 & \cdots & & 0 \\ 0 & w_2 & & & \\ \vdots & & 0 & & \\ & & & \ddots & \\ 0 & & & & 0 \end{pmatrix} \in \mathbb{R}^{6\times 6}. \tag{28}$$

To obtain the matrix of constraints $C$, we first rewrite the constraints to

$$l_1 - \tilde{y}_1 \leq 0 \tag{29}$$

$$\tilde{y}_1 - u_1 \leq 0 \tag{30}$$

$$l_2 - \tilde{y}_2 \leq 0 \tag{31}$$

$$\tilde{y}_2 - u_2 \leq 0 \tag{32}$$

$$u_1 - l_2 \leq 0, \tag{33}$$

which is equivalent to

$$\tilde{\xi}_3 - \tilde{\xi}_1 \leq 0 \tag{34}$$
$$\tilde{\xi}_1 - \tilde{\xi}_5 \leq 0 \tag{35}$$
$$\tilde{\xi}_4 - \tilde{\xi}_2 \leq 0 \tag{36}$$
$$\tilde{\xi}_2 - \tilde{\xi}_6 \leq 0 \tag{37}$$
$$\tilde{\xi}_5 - \tilde{\xi}_4 \leq 0. \tag{38}$$

These inequalities can be formulated to $C\tilde{\xi} \leq 0$ with

$$C = \begin{pmatrix} -1 & 0 & 1 & 0 & 0 & 0 \\ 1 & 0 & 0 & 0 & -1 & 0 \\ 0 & -1 & 0 & 1 & 0 & 0 \\ 0 & 1 & 0 & 0 & 0 & -1 \\ 0 & 0 & 0 & -1 & 1 & 0 \end{pmatrix}. \tag{39}$$

### 4 Implementation

Parameter estimations were run on a Intel(R) Xeon(R) Gold 6126 @ 2.60GHz processor with 384 GB RAM. For all models and methods, each of the local optimizations were run on a single core with a wall-time limit of 2h for model M4 and 1h for the other models. Profiles were run on a Intel(R) Core(TM) i5-6200U CPU @ 2.30GHz with 4 GB RAM. The gradient-based optimization was performed using the L-BFGS-B algorithm from SciPy (Jones *et al.*, 2001) with default options except for `fatol` $= 10^{-8}$ and `gtol` $= 10^{-8}$. The gradient-free optimization was run with SciPy's Powell algorithm with default options except for `fatol` $= 10^{-8}$.

For profile calculation, we used the implemented routines in pyPESTO. We adapted some hyperparameters, namely `min_step_size` $= 0.0001$, `delta_ratio_max` $= 0.00005$, `default_step_size` $= 0.0005$, `reg_order` $= 4$ and `reg_points` $= 50$. Additionally, we adapted the optimizer tolerances to `fatol` $= 10^{-10}$ and `gtol` $= 10^{-10}$.

For the inner optimization, we used the reparameterized and reduced formulation proposed by Schmiester *et al.* (2020). As optimizer, we chose SciPy's L-BFGS-B with `maxiter` $= 2000$ and `ftol` $= 10^{-10}$. The linear system of equations for gradient computation was solved using the sparse solver spsolve from `scipy.sparse.linalg`.

As weights, we used

$$w = \sum_i y(t_i, \theta) + \varepsilon, \quad \varepsilon = 10^{-8}. \tag{40}$$

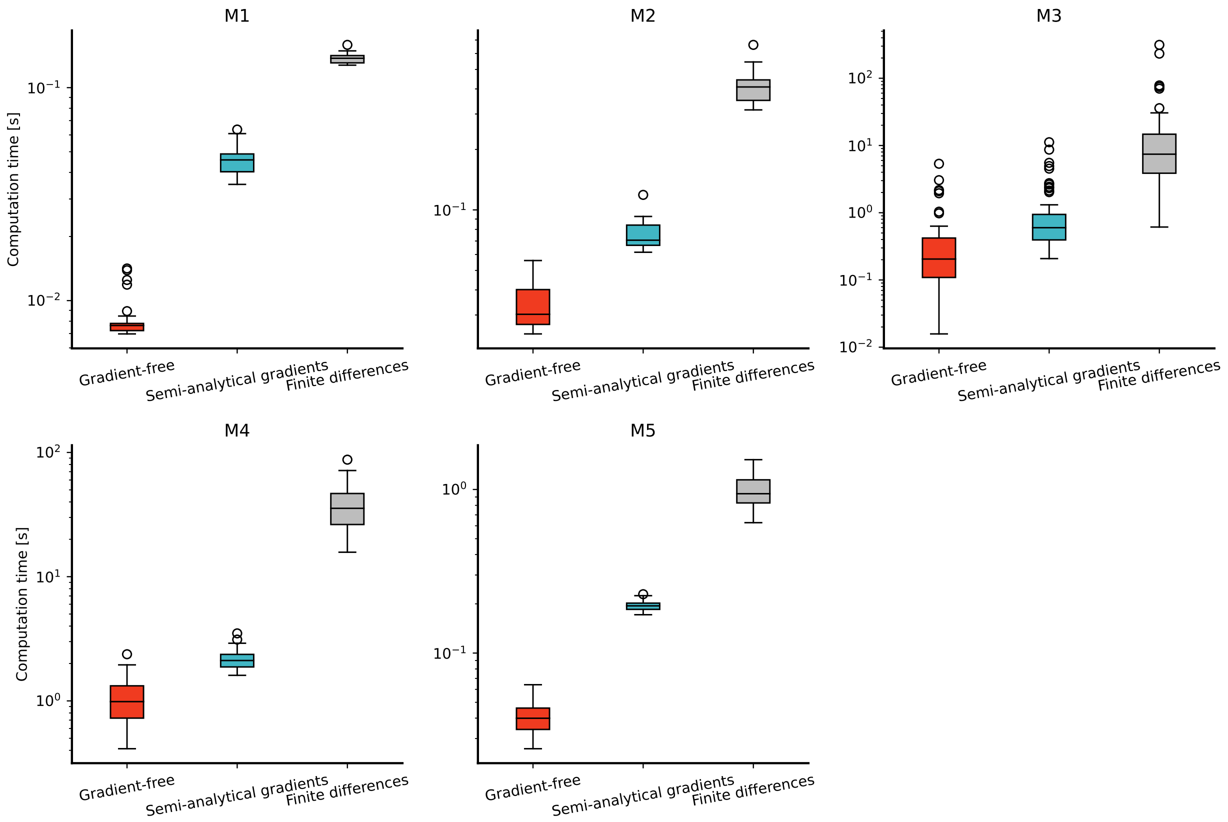


Figure S1: Computation times for an objective function evaluation for 50 randomly sampled parameter vectors for all five application examples without gradients, and with gradients using the semi-analytical approach and central finite differences.

As minimal values for the interval and gap sizes, we used

$$\begin{aligned} g &= \frac{\max(y(\theta))}{4(n_k-1)+1} + \gamma \\ s &= \frac{\max(y(\theta))}{2n_k+1} \end{aligned} \tag{41}$$

with $\gamma = 10^{-1}$ for models M2 and M4, and $\gamma = 10^{-2}$ for models M1, M3 and M5.

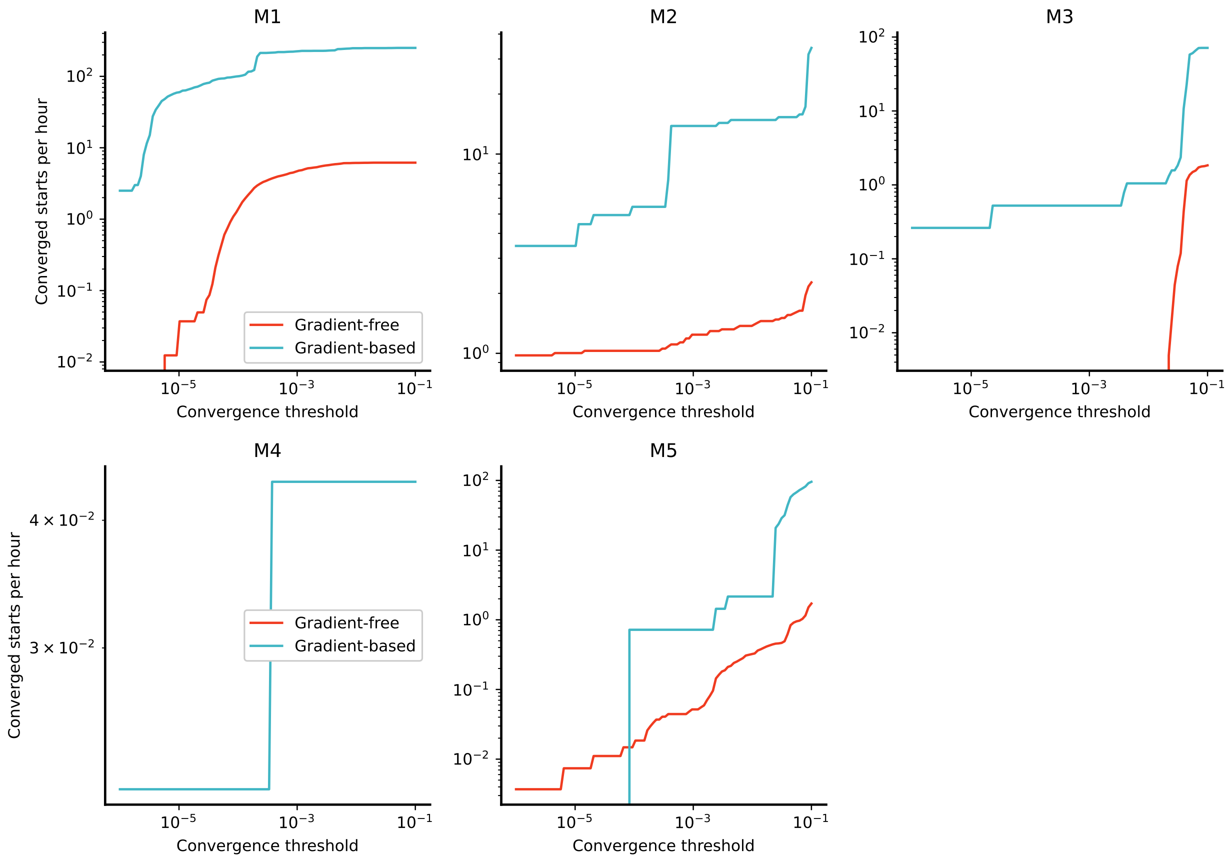


Figure S2: Number of converged starts per hour for different convergence thresholds, i.e. maximal absolute differences to the overall best objective function value.

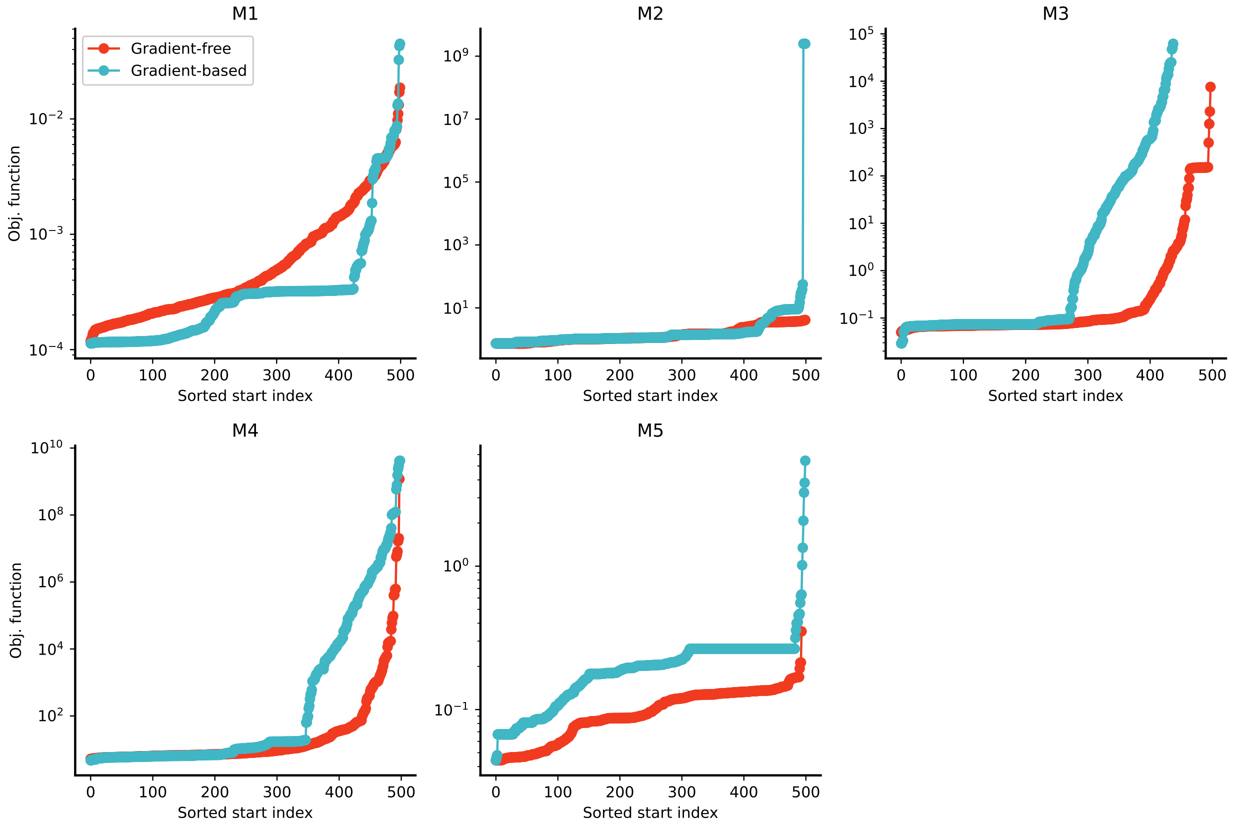


Figure S3: Waterfall plots for all models using gradient-free and gradient-based optimization. All 500 starts are shown.

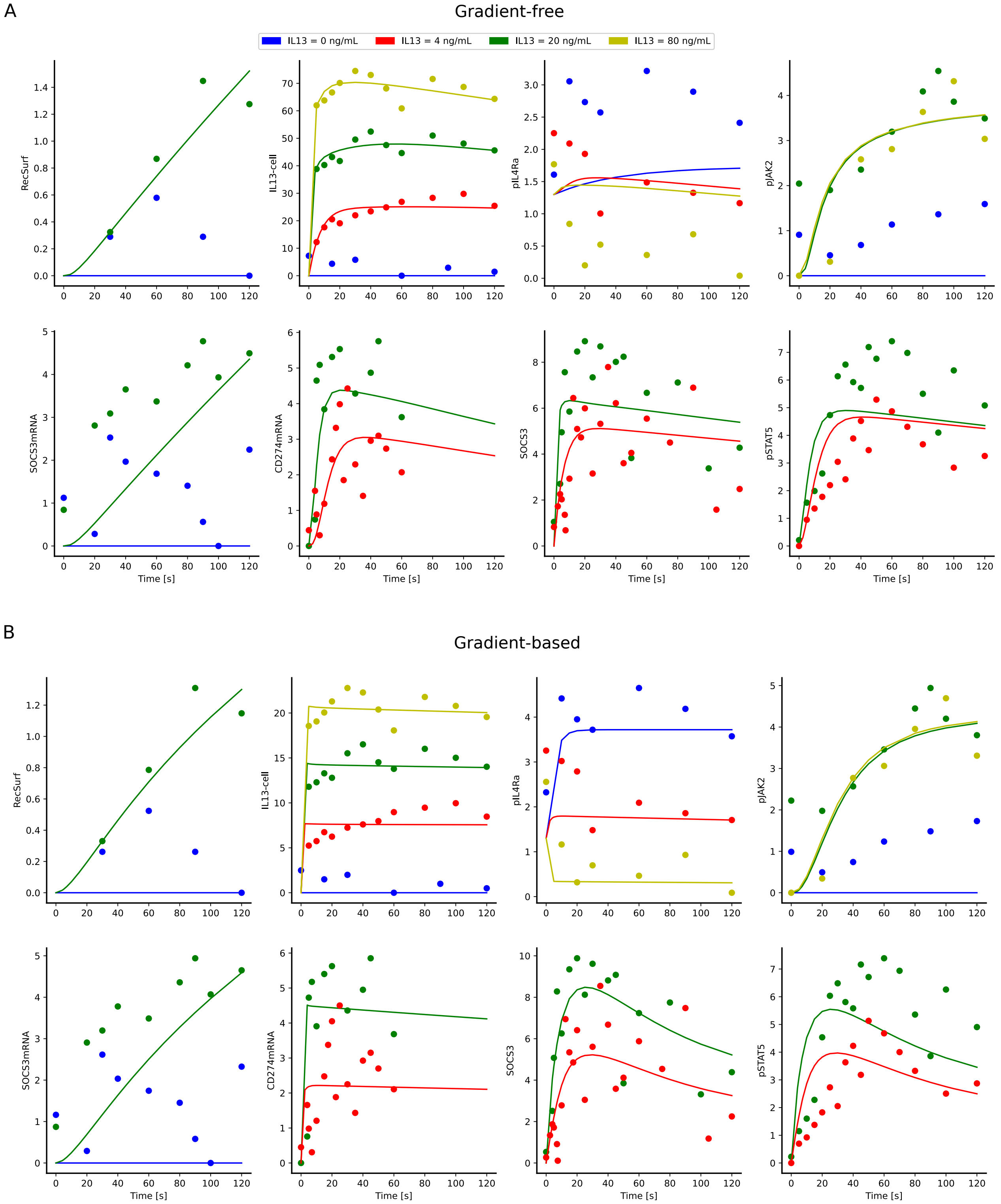


Figure S4: Model simulation and optimal surrogate data for the best found parameters of the gradient-free (A) and gradient-based (B) optimization for model M4.

Jones, E., Oliphant, T., Peterson, P., *et al.* (2001). SciPy: Open source scientific tools for Python.
